# Fungal effector SIB1 of *Colletotrichum orbiculare* has unique structural features and can suppress plant immunity in *Nicotiana benthamiana*

**DOI:** 10.1101/2021.07.01.450775

**Authors:** Ru Zhang, Noriyoshi Isozumi, Masashi Mori, Ryuta Okuta, Suthitar Singkaravanit-Ogawa, Tomohiro Imamura, Pamela Gan, Ken Shirasu, Shinya Ohki, Yoshitaka Takano

## Abstract

Functional screening of effector candidates using a transient expression assay in *Nicotiana benthamiana* identified two virulence-related effectors, named SIB1 and SIB2 (<u>S</u>uppression of <u>I</u>mmunity in *N. <u>b</u>enthamiana*), of an anthracnose fungus *Colletotrichum orbiculare*, which infects both cucurbits and *N. benthamiana. Agrobacterium*-mediated transient expression of SIB1 or SIB2 increased the susceptibility of *N. benthamiana* to *C. orbiculare*, which suggested these effectors can suppress immune responses in *N. benthamiana*. The presence of SIB1 and SIB2 homologs was found to be limited to the genus *Colletotrichum*. SIB1 suppressed both the generation of reactive oxygen species (ROS) triggered by the bacterial pathogen-associated molecular pattern (PAMP), flg22, and the cell death response triggered by the *Phytophthora infestans* INF1 elicitin in *N. benthamiana*. We determined the NMR-based structure of SIB1 to obtain its structural insights. The three-dimensional structure of SIB1 comprises five β-strands, each containing three disulfide bonds. The overall conformation was found to be a cylindrical shape, such as the well-known antiparallel β-barrel structure. However, the β-strands were found to display a unique topology, one pair of these β-strands formed a parallel β-sheet. These results suggest that the effector SIB1 present in *Colletotrichum* fungi has unique structural features and can suppress PAMP-triggered immunity (PTI) in *N. benthamiana*.

## Introduction

Plants use multilayered strategies to detect and defeat pathogenic microbes trying to attack them (1,2). As the first layer of plant defense, plants recognize conserved components of microbes called PAMPs, which are often present on their external face. Plant recognition of PAMPs triggers PTI. Although the plant immune system against most potential pathogenic microbes, especially nonadapted pathogens, is thought to depend mainly on PTI, adapted pathogens have evolved various mechanisms to suppress PTI (3). The secreted virulence factors, called effectors, play important roles in the suppression of PTI. In response to a pathogen’s use of effectors to try to suppress PTI, plants actuate their second layer of defense, called effector-triggered immunity (ETI) (4). ETI induces strong and robust immune responses that are typically associated with programmed cell death (PCD), a response referred to as the hypersensitive response (HR).

Members of the ascomycete genus *Colletotrichum* include numerous species that can infect a wide range of plant species, including many commercially important cultivars (5-7). The lifestyle of *Colletotrichum* species is considered to be hemibiotrophic, which combines an initial short biotrophic phase to maintain live host tissue and a subsequent necrotrophic phase that kills host tissue. In general, *Colletotrichum* fungi develop a specialized infection structure called appressorium that is darkly pigmented with melanin, and melanized appressorium is important for host penetration (8,9). Genome analyses have identified numerous effector candidate genes in *Colletotrichum* fungi such as *C. higginsianum* and *C. orbiculare* (5,6).

*C. orbiculare* belongs to the orbiculare clade and infects multiple cucurbitaceous cultivars (9-11). Interestingly, *C. orbiculare* can also infect *Nicotiana benthamiana*, which belongs to the Solanaceae family but is distant from cucurbits (12-14). We previously reported on the virulence-related effectors NIS1 and CoDN3 of *C. orbiculare* that are preferentially expressed in the biotrophic phase (15,16). We reported that the expression of NIS1 leads to PCD in *N. benthamiana*, and that the NIS1-triggered PCD is suppressed by CoDN3 expression (16). CoDN3 also inhibits PCD in *N. benthamiana* induced by another *C. orbiculare* effector NLP1 (17). We recently reported on the NIS1 targets *Arabidopsis thaliana* BAK1 and BIK1, which function in PAMP recognition and subsequent PTI activation, together with pattern recognition receptors that sense particular PAMPs (18).

We have previously reported that both adapted and nonadapted *Colletotrichum* fungi commonly develop melanized appressoria on *Arabidopsis* at 1 day post inoculation (1 dpi) (19). However, melanized appressoria of the adapted *Colletotrichum* fungus develop invasive hyphae successfully, whereas those of nonadapted *Colletotrichum* fungi fail to develop invasive hyphae because *Arabidopsis* plants activate a preinvasive defense (19). The finding therefore suggested that melanized appressoria of *Colletotrichum* fungi likely secrete effectors that are critical for the suppression of preinvasive plant defense. Consistently, microarray-based expression analysis of *C. orbiculare* inoculated on *N. benthamiana* shows that many small, secreted protein genes are highly expressed at 1 dpi, when the pathogen has developed melanized appressoria but has not yet formed invasive hyphae (5).

In this study, to identify novel virulence-related effectors of *C. orbiculare*, we focused on the effector-like genes expressed at 1 dpi after inoculation of *C. orbiculare* on *N. benthamiana*. Using the newly obtained RNA sequence data derived from *N. benthamiana* inoculated with *C. orbiculare* at 1 dpi, we selected candidate effector-like genes of *C. orbiculare* and subjected them to a functional screening assay to assess *Agrobacterium*-mediated transient expression. Each candidate was transiently expressed in *N. benthamiana* leaves that were subsequently challenged with *C. orbiculare* to assess each candidate’s ability to suppress the immunity of *N. benthamiana*. In these experiments, we identified two novel virulence-related effectors, named SIB1 and SIB2 (<u>S</u>uppression of <u>I</u>mmunity in *N. <u>b</u>enthamiana*), that suppressed *N. benthamiana* immunity against *C. orbiculare*. We then performed further characterization of SIB1. Transient expression of SIB1 suppressed both the generation of reactive oxygen species (ROS) triggered by a bacterial PAMP, flg22, and the cell death response triggered by the *Phytophthora infestans* INF1 elicitin. We next determined the tertiary structure of SIB1 to obtain structural insights into this effector. Using NMR analysis, we have solved the tertiary structure of SIB1, which showed that the effector SIB1 of *C. orbiculare* has unique structural features.

## Results

### Functional screening of virulence-related effectors in *C. orbiculare*

We obtained RNA sequence data from the following: (i) *N. benthamiana* leaves inoculated with *C. orbiculare* at 1, 3, and 7 dpi; (ii) conidia of *C. orbiculare*, and (iii) *in vitro* grown hyphae of *C. orbiculare*. We ranked the putative secreted protein genes of *C. orbiculare* based on their expression in *N. benthamiana* at 1 dpi (Table S1). The list included *NIS1* and *CAD1*, which we have previously studied (18,20). We then selected eight candidates, named CE1 to CE8 (Table S1), from the list, and these selected candidates were subjected to further functional screening. As mentioned above, *C. orbiculare* infects and causes lesions in *N. benthamiana* (13,14). In a study using a functional assay based on the *Agrobacterium*-mediated transient expression of NIS1 in *N. benthamiana* and subsequent inoculation with *C. orbiculare*, we recently reported that the expression of the effector NIS1 in *N. benthamiana* increased its susceptibility to *C. orbiculare* (18). We applied this assay to the functional screening of the selected candidates.

We expressed each candidate in *N. benthamiana* by transient expression using *Agrobacterium* infiltration, and we challenged the expression sites of each candidate by inoculation with *C. orbiculare*. The expression of CE6 caused lesion development before *C. orbiculare* inoculation, suggesting that CE6 can induce cell death in *N. benthamiana* (Fig. S1). The other candidates did not cause lesion development before *C. orbiculare* inoculation. Notably, we found that the expression of two candidates (CE7 and CE5) in *N. benthamiana* increased its susceptibility to *C. orbiculare* (Fig. 1A and 1B). Expression of the other candidates had no obvious effect on the susceptibility to *C. orbiculare*. These results suggest that CE7 and CE5 can suppress plant immunity of *N. benthamiana* against *C. orbiculare*.

**Figure 1.**
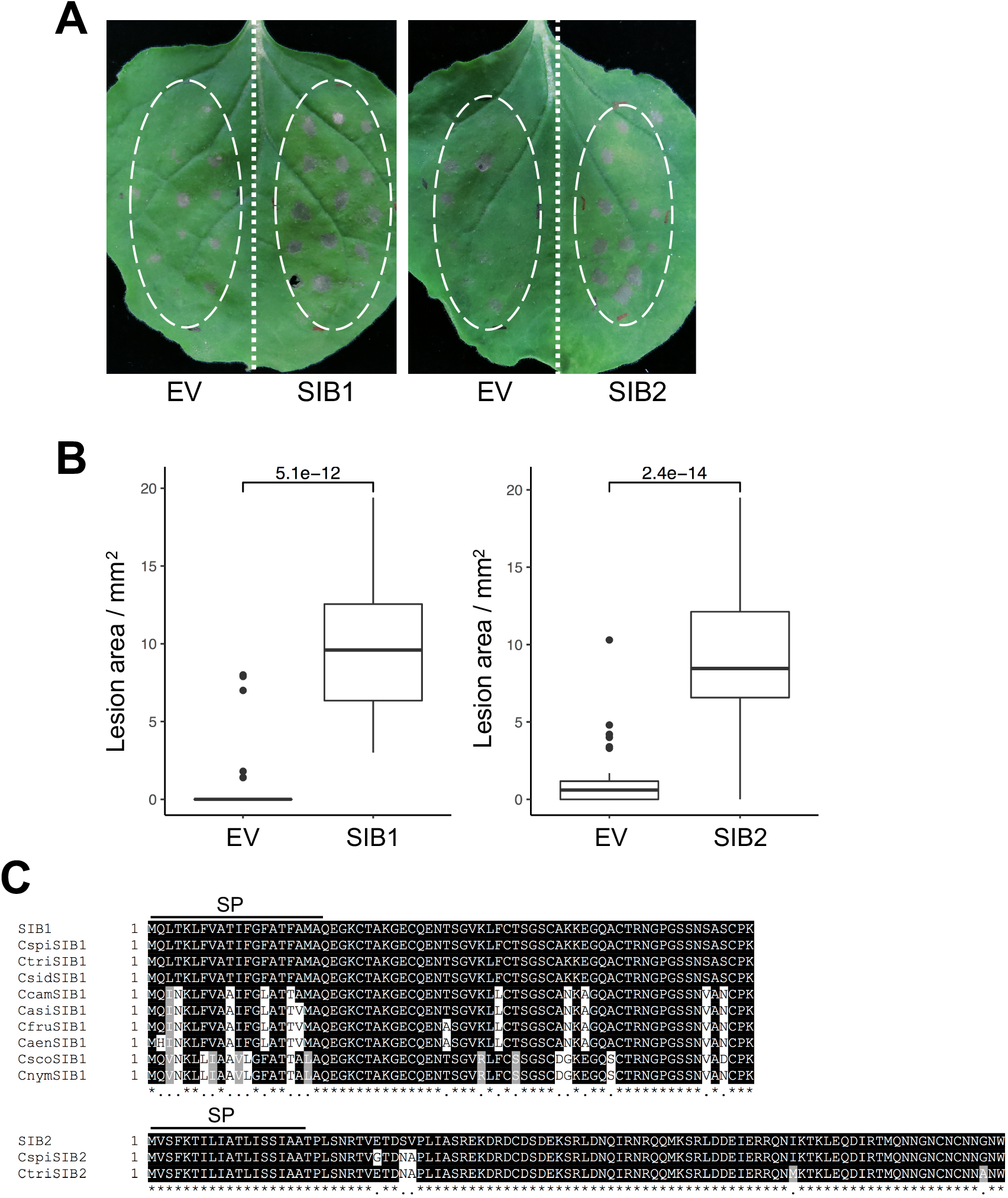
Decreased immunity to *C. orbiculare* caused by transient expression of the effector SIB1 or SIB2 in *N. benthamiana*. (A) Increased lesion development of *C. orbiculare* on *N. benthamiana* when *SIB1* or *SIB2* were transiently expressed. *N. benthamiana* leaves were infiltrated by *A. tumefaciens* harboring the plasmid expressing *SIB1*, the plasmid expressing *SIB2*, or the empty plasmid (EV), and the infiltrated leaves were incubated for 2 days and then drop-inoculated with conidial suspensions (5 ×10^5^ conidia/mL) of *C. orbiculare* 104-T. The photograph was taken at 5 dpi. (B) Quantification of lesion size on *N. benthamiana* leaves transiently expressing *SIB1* or *SIB2* after *C. orbiculare* inoculation. The lesion size in Fig. 1A was measured using ImageJ software for three biological replicates. The *t* test was used to identify significant differences. (C) *SIB1* and *SIB2* are conserved in *Colletotrichum* species. The amino acid sequence alignment of SIB1 or SIB2 with their orthologs of *Colletotrichum* species were shown. The alignments include the orthologs showing more than 75% amino acid identity obtained using a BlastP search of the NCBI non-redundant protein database using SIB1 or SIB2 as the query sequences. They were derived from the diverse *Colletotrichum* species represented by Cspi (*C. spinosum*), Ctri (*C. trifolii*), Csid (*C. sidae*), Ccam (*C. camelliae*), Casi (*C. asianum*), Cfru (*C. fructicola*), Caen (*C. aenigma*), Csco (*C. scovillei*), and Cnym (*C. nymphaeae*). The alignments were made using the ClustalW program. Identical residues in SIB1 or SIB2 are shaded in black, and conserved residues are shaded in gray. SP indicates the putative signal peptide region.

We named CE7 and CE5 as *SIB1* and *SIB2* (<u>S</u>uppression of <u>I</u>mmunity in *N. <u>b</u>enthamiana*), respectively. *SIB1* (GenBank accession number, TDZ19150.1) encodes a protein of 70 amino acids that had no significant matches in a Pfam search. SignalP analysis has suggested that SIB1 has a signal peptide of 20 amino acids (21) (Fig. 1C). *SIB2* (GenBank accession number, TDZ19243.1) encodes a protein of 99 amino acids that includes a signal peptide of 18 amino acids but has no clear domains as shown in a Pfam search. We then performed BlastP against the NCBI non-redundant protein database using SIB1 and SIB2 as the query sequences (Fig. S2). We found homologs of *SIB1* (100% identity in amino acid sequence) in *C. spinosum, C. trifolii*, and *C. sidae* that are members of the orbiculare clade that *C. orbiculare* belongs to. In contrast, genes predicted to encode full-length SIB2 homologs were only identified in *C. spinosum* and *C. trifolii*, but not in *C. sidae*. Homologs of both *SIB1* and *SIB2* were also found in a subset of *Colletotrichum* species outside the orbiculare clade but were not found outside the *Colletotrichum* genus (Fig. S2).

### Suppression by SIB1 of multiple PTI responses in *N. benthamiana*

We decided to focus on SIB1 and performed further characterization of this novel effector. We will report further studies on CE6 and SIB2 elsewhere. To investigate whether SIB1 suppresses PAMP-triggered ROS generation in *N. benthamiana*, we measured the ROS generation triggered by a bacterial PAMP flg22 in *N. benthamiana* expressing SIB1. It was recently reported that the NIS1 of *C. orbiculare* and an NIS1 homolog of *Magnaporthe oryzae* (MoNIS1) commonly suppress flg22-induced ROS production in *N. benthamiana* (18); therefore, we also investigated flg22-induced ROS production in the presence of MoNIS1. Both SIB1 and MoNIS1 suppressed flg22-induced ROS production compared with the negative control enhanced green fluorescent protein (eGFP) (Fig. 2A), which suggests that SIB1 can suppress one of the typical PTI responses. Some effectors have been shown to increase the virulence of a pathogen by suppressing the HR, which is accompanied by cell death (22).

**Figure 2.**
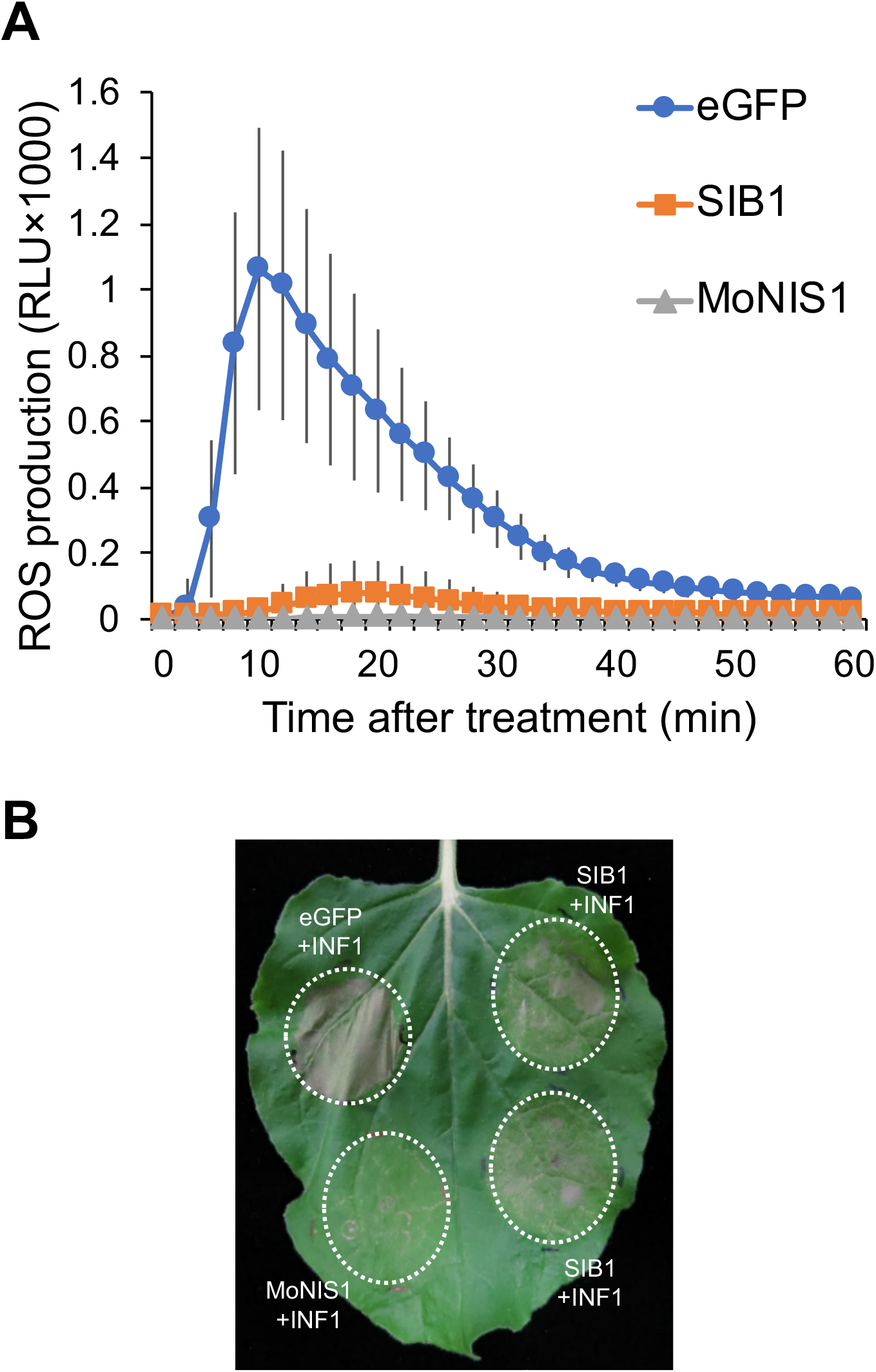
Suppression of *SIB1* by PAMP-triggered ROS generation and HR cell death in *N. benthamiana*. (A) flg22-triggered ROS production in *N. benthamiana* was inhibited by transient expression of *SIB1*. After treatment with 500 nM flg22, the total ROS production was measured in *N. benthamiana* transiently expressing *SIB1-HA*, or *MoNIS1-HA* (positive control), or *eGFP* (negative control). (B) Partial suppression of *INF1*-induced cell death by *SIB1. N. benthamiana* leaves were first infiltrated with *A. tumefaciens* harboring a plasmid expressing *SIB1, eGFP* (negative control), or *MoNIS1* (positive control). After 1 day, the second infiltration with *A. tumefaciens* harboring a plasmid expressing *INF1* was performed, and the infiltrated leaves were incubated for 5 days.

*P. infestans* INF1 is a well-known oomycete PAMP elicitor that can induce the HR in *N. benthamiana* leaves (23). NIS1 and MoNIS1 also suppress INF1-induced HR cell death in *N. benthamiana* (18). To investigate whether SIB1 can interfere with cell death triggered by the PAMP elicitor, SIB1, MoNIS1 or eGFP was expressed in *N. benthamiana* using *Agrobacterium* infiltration, and the infiltration sites were challenged with *Agrobacterium* carrying *INF1*. INF1-triggered lesion development was observed in the infiltration sites expressing GFP but was clearly suppressed in the sites expressing MoNIS1 as previously shown. Notably, SIB1 also suppressed INF1-induced lesion development, which indicated that SIB1 suppresses HR cell death triggered by the PAMP elicitor INF1 (Fig. 2B). These findings suggest that the effector SIB1 can suppress multiple PTI responses in *N. benthamiana*.

We next performed quantitative reverse transcription PCR (RT-qPCR) analysis to investigate the expression pattern of *SIB1* in conidia of *C. orbiculare* inoculated on *N. benthamiana* and cucumber. The expression of *SIB1* at 0, 24, and 72 hour post inoculation (hpi) with *C. orbiculare* on *N. benthamiana* was consistent with the RNA sequence data (conidia, 1 dpi in *Nb*, and 3 dpi in *Nb*) (Table S1). *SIB1* expression started to be induced at 8 hpi and its expression level was highest at 12 hpi (Fig. 3A). *SIB1* expression was induced after inoculation on cucumber (Fig. 3A). However, the expression pattern of *SIB1* on cucumber was not identical to that on *N. benthamiana* (Fig. 3A); for example, *SIB1* expression was highly induced at 72 hpi on cucumber, but not on *N. benthamiana*.

**Figure 3.**
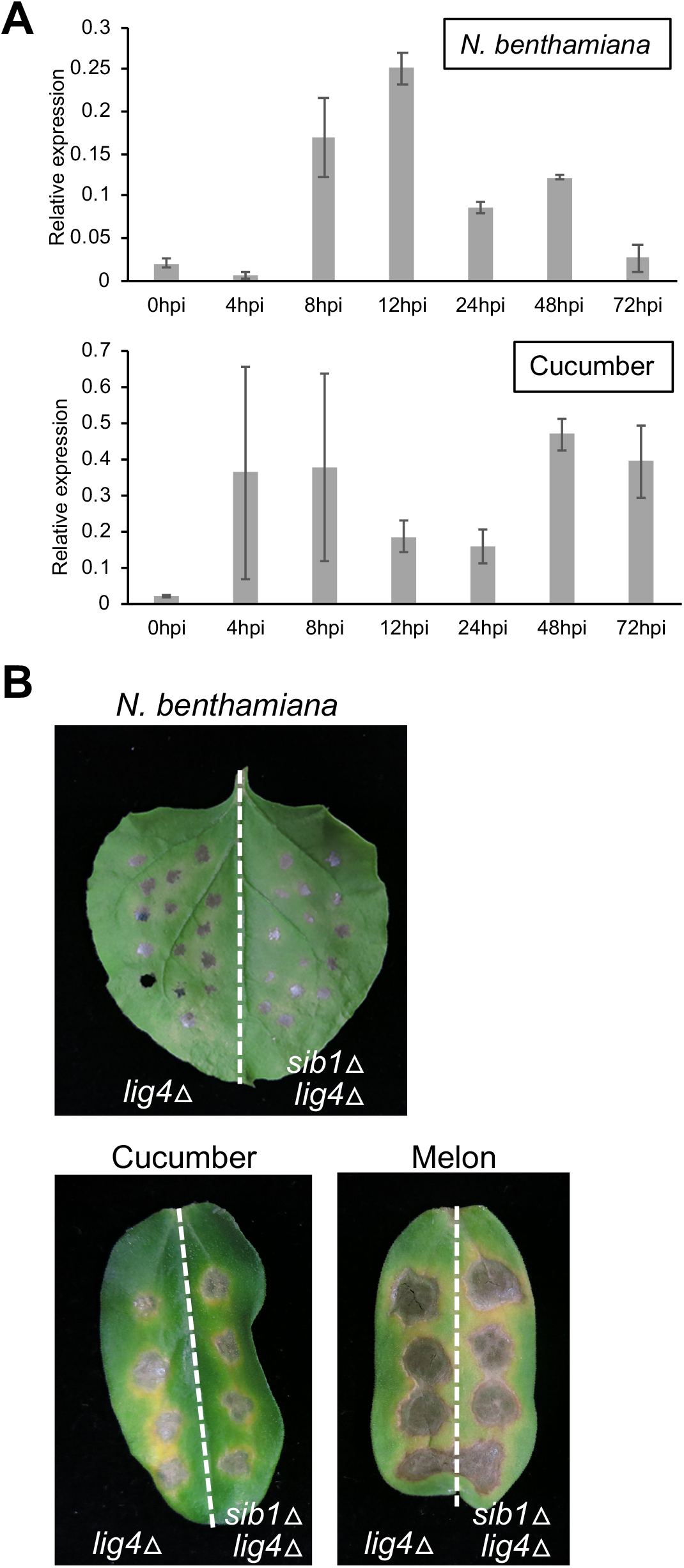
Gene expression and knockout analysis of *SIB1*. (A) Expression pattern of *SIB1* in *C. orbiculare* inoculated on *N. benthamiana* and cucumber. The conidial suspension of *C. orbiculare* wild-type strain (1 × 10^6^ conidia per milliliter) was inoculated on *N. benthamiana* leaves or cucumber cotyledons. The total RNA of inoculated plants was extracted and subjected to RT-qPCR analysis to investigate *SIB1* expression. The *C. orbiculare* actin gene was used as the internal control. Mean and SD were calculated from three independent samples. (B) Gene disruption of *SIB1* had no visible effects on the virulence of *C. orbiculare* inoculated on *N. benthamiana*, cucumber, or melon. Conidial suspension (5 × 10^5^ conidia/mL) of the parental *lig4*Δ strain or the *sib1*Δ strain (*lig4*Δ background) was drop-inoculated on *N. benthamiana* leaves, cucumber cotyledons, and melon cotyledons, and the inoculated plants were incubated at 24°C for 7 days.

We next applied targeted gene disruption of *SIB1* and investigated whether *SIB1* is required for the virulence of *C. orbiculare*. To delete *SIB1* in *C. orbiculare*, we first generated the *lig4*Δ strain from *C. orbiculare* 104-T, in which an increased homologous recombination ratio is expected (24), and used the *lig4*Δ strain as the parental strain for the gene disruption of *SIB1* (<u>details are included in Materials and Methods</u>). The *SIB1*-knockout vector, named pCB1636SIB1, was constructed and introduced into the *lig4*Δ strain, and knockout mutants of *SIB1* were obtained (Fig. S3A and 3B). The colony morphology and conidiogenesis of the generated *SIB1*-knockout mutants (*sib1*Δ) on potato dextrose agar (PDA) medium were similar to those of the control parental strain (Fig. S3C). We then inoculated the *sib1*Δ strains on *N. benthamiana*, cucumber, and melon, and found that the *sib1*Δ strains developed the same lesions as the control strain for all plants tested (Fig. 3B).

### Posttranscriptional modification of SIB1

We next focused on the structural aspects of the effector SIB1. We tried to produce SIB1 protein in the suspension-cultured BY-2 system (25,26), in which the research target protein is expressed as a fused protein together with both tobamovirus (ToMV) and transcription factor (XVE) to increase the productivity. This system also uses optimized signal peptides for endoplasmic reticulum migration and secretion to fold the yielded protein. The system can produce proteins containing disulfide bonds in their native conformation (27-30). We used this system to prepare SIB1, whose amino acid sequence has six Cys residues that are expected to form intramolecular disulfide bonds.

We prepared semi-purified SIB1 protein. When all Cys residues are in reduced form, the theoretical mass of SIB1 is calculated as 5414.19 *m/z*. We treated the purified SIB1 as for the reduced form and confirmed the mass. As shown in Fig. 4, the mass of the reduced SIB1 was slightly smaller than the theoretical value of 5396.388 *m/z*, which suggests that the SIB1 expressed by the BY-2 system had some modification. A search of the Unimod database suggested that the difference (–17.802 *m/z*) is derived from pyroglutamylation of the N-terminal residue, Gln1. To confirm the pyroglutamylation of SIB1, we used pyroglutamate aminopeptidase (PGAP) treatment of SIB1. Because only N-terminal pyroglutamic acid is cleaved by this treatment, we used this assay to determine whether the sample protein contained N-terminal pyroglutamic acid. For the PGAP-treated sample (lower panel of Fig. 4), only the peak (5267.895 *m/z*), which corresponds to the N-terminal glutamine-cleaved SIB1 (ΔQ1-SIB1), was detected. The mass spectrometry (MS) results showed clearly that the N-terminal residue of SIB1 expressed in BY-2 cells was pyroglutamic acid. The Gln at the N-terminal end was easily modified to pyroglutamic acid (31,32).

**Figure 4.**
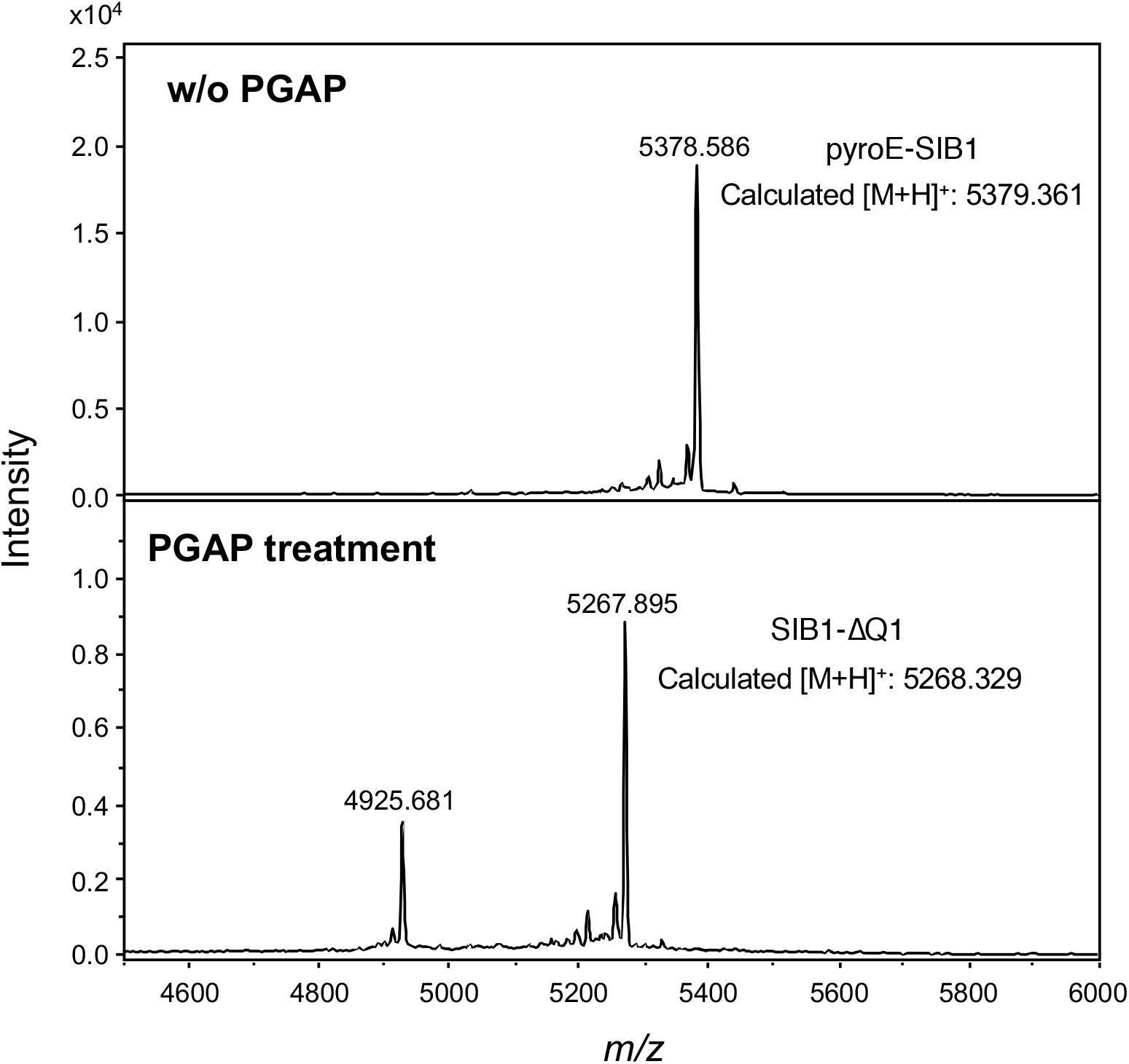
Confirmation of pyroglutamylation of SIB1. Pyroglutamate aminopeptidase (PGAP)-untreated (*upper panel*) and -treated (*lower panel*) SIB1 were analyzed using MALDI-TOF-MS.

The mass of pyroglutamylated SIB1 suggested that all Cys residues were in the oxidized form, as shown in Fig. 5A. Therefore, we used MS to analyze the disulfide bond pairs. Lys-C treatment under nonreducing conditions caused SIB1 digestion, but the disulfide bonds were maintained. As shown in Fig. 5B, we observed four peaks for the Lys-C-treated sample: one peak (5031.432 *m/z*) corresponding to undigested SIB1 and three other peaks (1014.473 *m/z*, 1949.780 *m/z*, and 2077.858 *m/z*) indicating digested peptides containing disulfide linkages. Further MS analysis of the products of the enzymatic digestion clearly indicated the existence of two disulfide bonds, Cys22–Cys27 and Cys35–Cys48, as shown in Fig. 5B. Peptides containing Cys5 and Cys11 were not detected, probably because of difficulty with their ionization. Because all Cys residues were in the oxidized form, as shown in Fig. 5B, the remaining two Cys residues, Cys5 and Cys11, were expected to form disulfide bonds.

**Figure 5.**
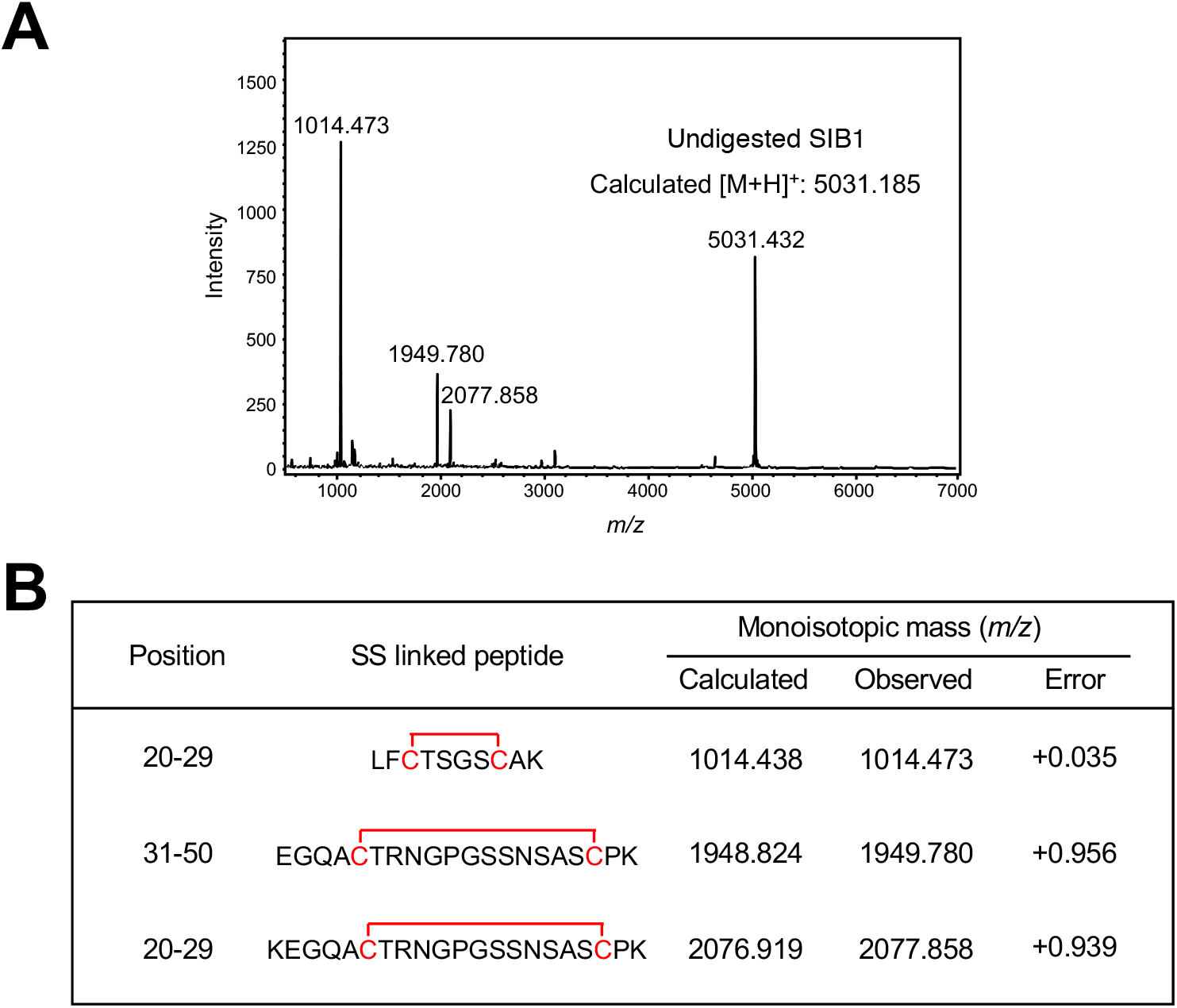
Determination of disulfide linkages of SIB1. (A) MALDI–TOF–MS spectra of SIB1 after Lys-C treatment. (B) The assignments of SS linked peptides of SIB1 obtained by Lys-C digestion.

### Structure of SIB1

Unlabeled and ^15^N-labeled SIB1 samples were expressed in the BY-2 system, and the N-terminus of NMR sample used in this study was pyroglutamylated. As shown in Fig. S4, the ^1^H-^15^N heteronuclear single quantum coherence (HSQC) spectrum of SIB1 showed well-dispersed signals with sharp line shapes, which indicated that SIB1 was in a stable conformation in solution. After the resonance assignments, three-dimensional structure calculation was performed using a standard CYANA protocol with the distance and angle constraints derived from NMR data. Disulfide bond constraints for three Cys–Cys pairs were also used in the calculation. We obtained the final 20 structures of SIB1 with 0.71 ± 0.07Å of root mean square deviation for all heavy atoms of residues 2 to 49. The structural statistics are summarized in Table 1. Although only 48% (24/50) of the residues formed secondary structure, no dihedral angle fell into the disallowed region in the Ramachandran plot. The final 20 structures showed no violation in distance or angle restraints.

**Table 1.**
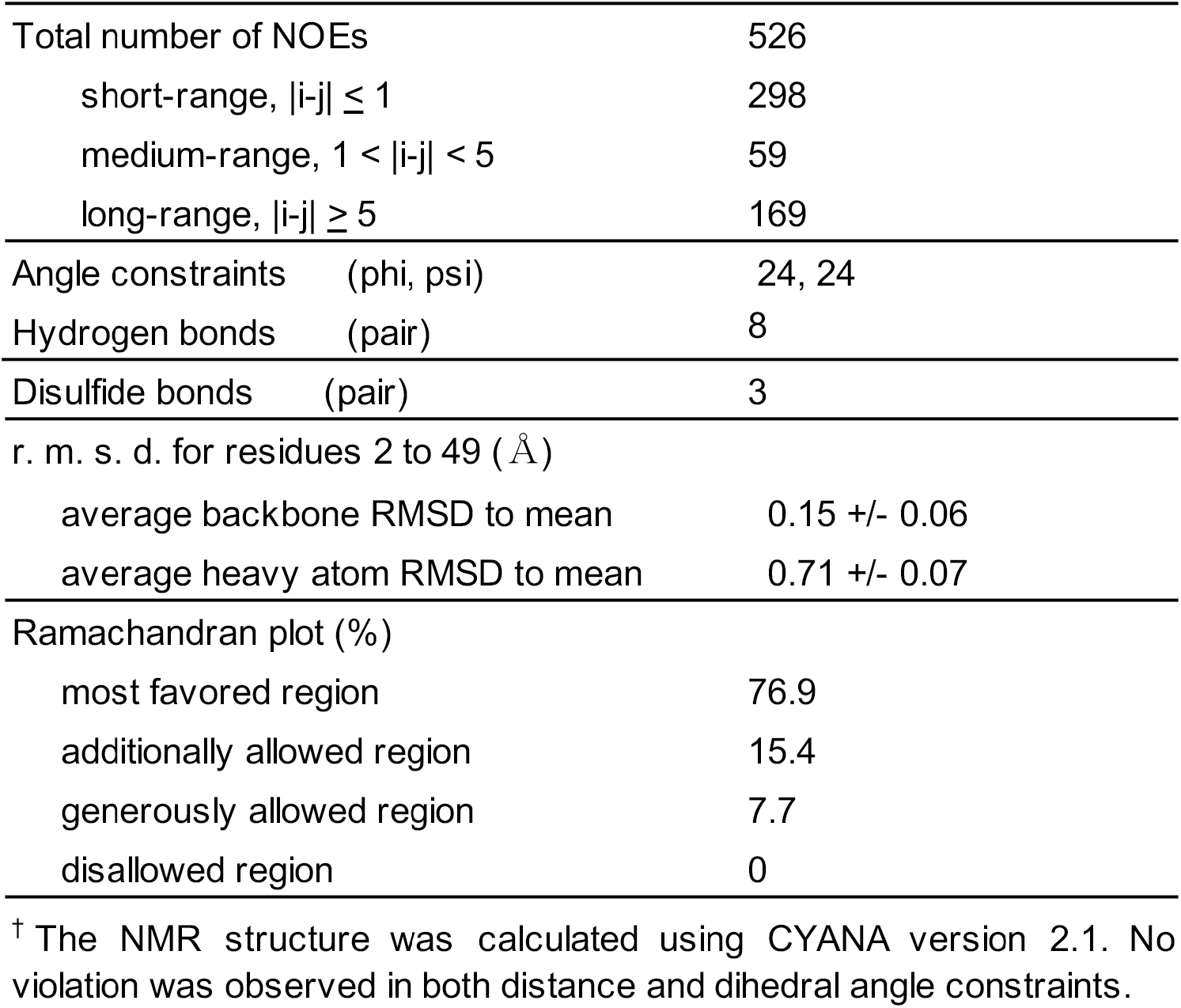
Statistics of the NMR structure calculation^†^

A backbone wire model of the final ensemble and a ribbon model of the representative model are shown in Fig. 6A and C, respectively. The three-dimensional structure of SIB1 comprised five β-strands without an α-helix. The strands form a cylindrical shape, the so-called β-barrel. The five strands were named β1 to β5 starting from the N-terminus span residue 2 to 6, 10 to 14, 18 to 23, 36 to 38, and 44 to 48, respectively. The topology of the five β-strands is shown in Fig. 6B. Although three pairs of the β-strands (β1–β2, β2– β3, and β4–β5) are in the antiparallel orientation, only one pair, β3–β5, adopts a parallel form. This is a unique characteristic of SIB1 because the antiparallel β-barrel is the most common structure. A search using the structure comparison server DALI (http://ekhidna2.biocenter.helsinki.fi/dali/) suggested that no protein in the database displays the SIB1-like five-strand β-barrel structure containing one parallel β-sheet. We found that the three-dimensional structure of SIB1 includes three disulfide bonds, all of which are located in the inner part of the molecule, as shown in Fig. 6C.

**Figure 6.**
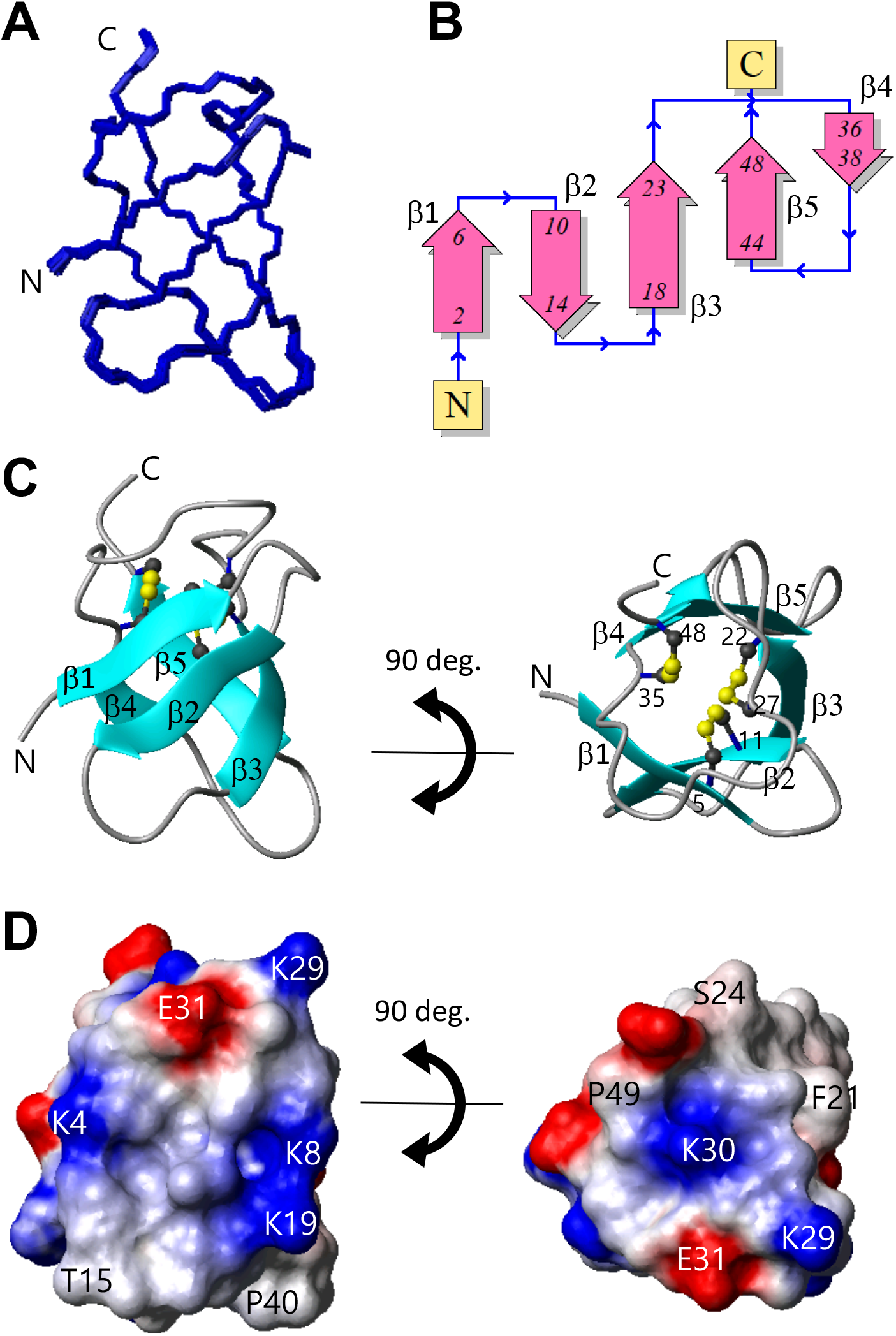
Three-dimensional structure of SIB1. Overlay of 20 NMR structures (A), topology of five β-strands (B), ribbon model of the representative structure using a ball-and-stick representation of the three disulfide bonds (C), and molecular surface charge distribution (D). Fig. 6B was generated on the PDB sum website (http://www.ebi.ac.uk/thornton-srv/databases/cgi-bin/pdbsum/GetPage.pl?pdbcode=index.html). The molecular surface shown in Fig. 6D is colored red (negative), blue (positive), and white (hydrophobic).

The electrostatic potential of the molecular surface of SIB1 is shown in Fig. 6D. An intriguing molecular surface property is seen at the top site of the β-barrel structure. A shallow bowl-like shape is formed by the long loop between β3 and β4. The central bed region of this area is positively charged, and this charge is surrounded by a hydrophobic rim.

As an additional analysis, we performed *T*_1_, *T*_2_, and {^1^H}-^15^N nuclear Overhauser effect spectroscopy (NOE) experiments to obtain information about the dynamics of each residue. These results are shown in Fig. S5. All 1/*T*_1_, 1/*T*_2_, and {^1^H}-^15^N NOE values indicated that the overall structure was rigid and stable in the NMR time scale. Slightly smaller {^1^H}-^15^N NOE values were observed only for the loop regions, which suggested that the loops are more flexible than the β-strand regions. Unlike the loop regions, relatively higher 1/*T*_2_ values were observed for few residues located in the β-strands, but such residues appeared sporadically through the amino acid sequence. Moreover, the 1/*T*_1_ and heteronuclear NOE did not show higher/lower values for such residues, indicating no further information about the rigidity. The analyses of the dynamics suggested that the poor plasticity of the SIB1 conformation makes it difficult to deduce the functional site involved in the conformational selection needed to adapt to the target. The classical key-and-lock binding manner might be proposed, but identification of the target binding region is not possible at present.

## Discussion

In this study, we selected effector candidate genes of *C. orbiculare* that were highly expressed at 1 dpi after inoculation of the pathogen on *N. benthamiana* and performed functional screening using an *Agrobacterium*-mediated transient expression assay in *N. benthamiana*. We identified CE6 as a factor that caused cell death in *N. benthamiana*. Importantly, we also identified two novel effectors of *C. orbiculare*, named SIB1 and SIB2, that suppressed *N. benthamiana* immunity against *C. orbiculare. SIB1* was found to be conserved in the genome of 18 *Colletotrichum* species but was not found outside the *Colletotrichum* genus. Homologs encoding the amino acid sequence identical to that of *C. orbiculare* SIB1 were found in *C. spinosum, C. trifolii*, and *C. sidae*, which also belong to the orbiculare clade (33). *SIB2* homologs were identified in the genome of 24 *Colletotrichum* species, including *C. spinosum* and *C. trifolii* but not in *C. sidae*.

We focused on SIB1 in this study. SIB1 suppressed the flg22-triggered ROS burst in *N. benthamiana*. Plant NADPH oxidases, also known as respiratory burst oxidase homologs (RBOHs), produce ROS (34). An RBOHB (NbRBOHB) of *N. benthamiana* plays crucial roles in ROS production triggered by PAMPs, such as bacterial flagellin and fungal chitin, and facilitates plant immunity against biotrophic pathogens such as the oomycete pathogen *P. infestans* (35-37). SIB1 also partially suppressed INF1-induced cell death in *N. benthamiana*. It has been reported that the silencing of *Rboh* genes leads to a reduction and delay in HR cell death caused by INF1 in *N. benthamiana* (37). Therefore, SIB1-mediated suppression of the ROS burst may be involved in the SIB1-mediated suppression of INF1-induced cell death. On the other hand, *NbRbohB* silencing decreases resistance to *P. infestans* but not to *C. orbiculare* (35). Therefore, the increased susceptibility of *N. benthamiana* to *C. orbiculare* via transient expression of SIB1 is unlikely to depend on the SIB1-mediated suppression of the ROS burst. SIB1 may be able to suppress other immune responses in addition to the ROS burst.

In the case of *C. orbiculare* inoculation on *N. benthamiana*, RT-qPCR analysis suggested that the expression of *SIB1* was highest at 12 hpi, when the pathogen has already developed appressoria for host invasion, and was strongly reduced at 72 hpi. This result suggests that SIB1 may contribute to the primary stage of host invasion. By contrast, in the case of *C. orbiculare* inoculation on cucumber, the expression of *SIB1* was highest at 48 hpi and its expression level remained high at 72 hpi. These findings suggest that *C. orbiculare* changes the expression pattern of effector genes, including *SIB1*, during infection of two unrelated susceptible plants, cucumber and *N. benthamiana*. In addition, the inoculation assays using the *SIB1-*knockout mutants revealed that *SIB1* was not essential for the virulence of *C. orbiculare* on *N. benthamiana*, cucumber, and melon, although the transient expression of SIB1 in *N. benthamiana* increased the susceptibility to *C. orbiculare*. We now consider that other effectors of *C. orbiculare* may have functional redundancy with SIB1.

The three-dimensional structure of SIB1 comprises five β-strands each with three disulfide bonds. A pair of β-strands forms a parallel β-sheet and the others are antiparallel. We tried homology searches to find proteins with SIB1-like topology. A search of SAS (http://www.ebi.ac.uk/thornton-srv/databases/sas/) and 3D-BLAST (http://3d-blast.life.nctu.edu.tw/) found no similar structures in these databases. We also tried ProFunc (http://www.ebi.ac.uk/thornton-srv/databases/cgi-bin/profunc), and the survey suggested two antifungal proteins comprising five β-strands (PDB codes: 2KCN and 1AFP) as the structural neighbors, but both of them show all antiparallel β-barrel topology. These results suggest that SIB1 displays a novel structural feature and that these structural characteristics may be related to the functional mechanism of SIB1.

We observed pyroglutamylation of SIB1 at the N-terminus in the present study. Similar N-terminal modification has been reported for many peptides and proteins. For example, brazzein, a sweet-tasting protein of African plants that adopts a well-known protein fold seen in defensins and arthropod toxins, has a pyroglutamylated N-terminal end (38). This modification may be necessary for preventing protein degradation in host cells (39). Therefore, it is possible that the N-terminal modification of SIB1 occurs in nature and functions to extend its lifetime in plant cells. For further understanding of the molecular function of the effector SIB1, especially in the suppression of immunity in *N. benthamiana*, further studies are needed for the comprehensive mutational analyses of SIB1 based on the unraveled SIB1 structure and identification of the *N. benthamiana* proteins targeted by SIB1.

## Materials and Methods

### Fungal strains and culture condition

*C. orbiculare* strain 104-T (MAFF240422) (stock culture of the Laboratory of Plant Pathology, Kyoto University) was used as the wild-type strain. For targeted gene disruption of *SIB1*, we generated the *lig4*Δ strain from 104-T and used this strain as the parental strain in this study. All fungal strains were maintained on PDA medium (3.9% [wt/vol] PDA; Nissui, Tokyo, for 104-T) at 24°C in the dark.

### Plasmid constructions

To express candidate genes in plants, pBICP35-CE1–CE8 transient expression vectors under the control of the 35S promoter were constructed using an In-Fusion system (Clontech, TaKaRa). The fragment containing the cDNA of CE1 was amplified with the primers 35S_CE1_Fw and 35S_CE1_Rv. The fragment was contained in a *Bam*HI site and was introduced into the *Bam*HI site of pBICP35, producing pBICP35-CE1. The other candidate gene plasmids used for transient expression were constructed in a similar way as pBICP35-CE1.

To delete *LIG4* of *C. orbiculare* (GenBank accession number, TDZ18841), we first generated pBATTEFPGEN. The geneticin-resistant gene cassette was amplified from pII99 (40) with the primers GENAS1B and GENS1X, and the amplified fragment was digested with *Xba*I and *Bam*HI, and then introduced into pBATTEFP (41), resulting in pBATTEFPGEN. The 5′-upstream region of *LIG4* in *C. orbiculare* was amplified using genomic PCR with the primers CoLIG5SN2 and CoLIG5ASN2. The fragment was digested with *Not*I and introduced into pBATTEFPGEN, resulting in pBATTEFPGEN5L. The 3′-downstream region of *LIG4* was amplified with the primers CoLIG3SA5 and CoLIG3ASA5. The fragment was digested with *Apa*I and introduced into pBATTEFPGEN5L, resulting in pBATTEFPGENLIG4KO.

To delete *SIB1* of *C. orbiculare*, we constructed a gene-disruption vector, pCB1636SIB1, using the two-step In-Fusion strategy (Clontech, TaKaRa). First, the ∼2.0-kb upstream region of *SIB1* was amplified using PCR with the primers SIB1_Up_Fw and SIB1_Up_Rv, and the fragment was digested with *Apa*I. This fragment was then introduced into the *Apa*I-digested pCB1636 (42), resulting in pCB1636S5. Second, the ∼2.0-kb downstream region of *SIB1* was amplified using PCR with the primers SIB1_Down_Fw and SIB1_Down_Rv, and the fragment was digested with *Eco*RI. This fragment was introduced into the *Eco*RI-digested pCB1636S5, resulting in pCB1636SIB1. The primers used for plasmid construction are listed in Table S2.

To produce SIB1 protein in tobacco BY-2 cells, we designed the amino acid sequence for the SIB1 protein fused with an extracellular signal peptide of *Arabidopsis* chitinase (SP-SIB1). Next, artificial *SP-SIB1* was synthesized by optimizing the codons in tobacco and introducing restriction enzyme sites for cloning at both ends (IDT, Coralville, IA, USA; Table S2). The artificial *SP-SIB1* was introduced into a chemically inducible tobamovirus vector (pBICLBSER-ToMV) (28). The resultant plasmid was named pBICLBSER-ToMV-SP-SIB1.

### RNA isolation and sequencing

RNA was isolated as previously described (5). In brief, total RNA from conidia containing 3-day-old hyphae grown in potato dextrose broth at 25°C and infected *N. benthamiana* leaves at dpi 1, 3, and 7 were isolated using a Plant RNeasy Mini kit with DNase I treatment (Qiagen, Hilden, Germany). Three biological replicates were prepared for each tissue type. Unstranded RNAseq libraries were prepared from Poly(A)+-tailed RNA using a TruSeq Sample Prep kit according to the manufacturer’s instructions before sequencing on an Illumina HiSeq 2000 sequencer to 50 bp in single-read mode. Reads were mapped to the *C. orbiculare* genome (version 2 accession number, AMCV02000000) using STAR version 2.6.0a (43) with the setting --alignIntronMax 1000. Read counts were obtained using Rsubread (v1.32.2) (44) using the following settings: isGTFAnnotationFile = TRUE, GTF.featureType = “exon”, GTF.attrType = “Parent”. Reads per kilobase million values (45) were calculated using edgeR (46) after applying calcNormFactors.

### Agrobacterium tumefaciens-mediated transient expression assay in N. benthamiana

For the agroinfiltration assay, *N. benthamiana* plants (5 to 6 weeks old) were used. Plants were grown in a controlled environment chamber at 25°C with 16 h of illumination per day. Each construct was transformed into *Agrobacterium tumefaciens* strain GV3101 by electroporation. Each *Agrobacterium* was cultured in Luria–Bertani medium broth containing kanamycin (50 μg/mL), rifampicin (50 μg/mL), and gentamicin (50 μg/mL). The cells were harvested by centrifugation and then resuspended in MMA induction buffer (1 L of MMA: 5 g of Murashige and Skoog salts, 1.95 g of MES, 20 g of sucrose, and 200 μM acetosyringone, pH 5.6). All suspensions (OD600 = 0.3) of the *Agrobacterium* strains were incubated for 1 h before being infiltrated into *N. benthamiana* leaves using a needleless syringe.

### Virulence-enhancement assay

*N. benthamiana* leaves were infiltrated with each *A. tumefaciens*. The infiltrated leaves were incubated for 2 days, after which 10 μL of conidial suspensions (5 × 10^5^ conidia/mL) of the *C. orbiculare* wild-type strain were drop-inoculated onto the infiltration areas of detached *N. benthamiana* leaves. Inoculated leaves were incubated at 24°C for 5 days. Quantitative assessment of lesion development was obtained using ImageJ for three biological replicates

### Suppression assay of INF1-induced cell death

Each tested gene was expressed in the *A. tumefaciens*-mediated transient expression assay as mentioned above. At 1 day after the first agroinfiltration, the second agroinfiltration with recombinant *A. tumefaciens* carrying p35S-INF1 was performed at same infiltration site. All suspensions (OD600 = 0.3) of the *Agrobacterium* strains were incubated for 1 h before infiltration. The suspensions were infiltrated into *N. benthamiana* leaves using a needleless syringe. INF1-induced lesions were observed at 3 to 5 days after the second infiltration.

### ROS assay

ROS production was monitored using a luminol-based assay (47). Leaf discs were made using a circular borer (diameter, 5 mm), and the collected leaf discs were incubated overnight in distilled water. For measurement of ROS production, leaf discs were placed in a 96-well plate containing 50 μL of distilled water and 50 μL of assay solution containing 20 mM Luminol (Sigma-Aldrich, St. Louis, A8511), and 1 mg/mL peroxidase (Sigma-Aldrich, St. Louis, P6782) and 0.5 μM flg22 (Invitrogen) were added to the wells. Luminescence was measured using a Luminoskan Ascent 2.1 (Thermo Fisher Scientific, Yokohama, Japan).

### RT-qPCR analysis of *SIB1* expression

Cucumber cotyledons were drop-inoculated with conidial suspension (1 × 10^6^ conidia/mL) of the *C. orbiculare* wild-type strain covering as much as possible of the abaxial surface. After incubation for 0, 4, 8, 12, 24, 48, and 72 h, the inoculated epidermis containing the fungal cells was peeled off from three cotyledons for each sample and immediately frozen in liquid nitrogen to fix the gene expression profile. As for the preparation of 0 h samples, once conidial suspensions were inoculated, inoculated epidermis were immediately peeled off. As for inoculation on *N. benthamiana*, leaves were spray-inoculated with conidial suspension (1 × 10^6^ conidia/mL) of the *C. orbiculare* wild-type strain. Then the whole leaves were frozen at particular time point in liquid nitrogen to fix gene expression profiles, one leaf for each sample. The frozen tissues were ground and total RNA was extracted by using the Agilent Plant RNA Isolation Mini Kit (Agilent Technologies). Three biological replicates were prepared for each time point. The relative gene expression of *SIB1* was assessed by RT-qPCR using primers SIB1_qRT_F and SIB1_qRT_R (Table S2). The TB Green™Premix Ex Taq™(TaKaRa) was used with a Thermal Cycler Dice Real Time System TP800 (TaKaRa) for RT-qPCR. The relative expression levels were normalized against the *C. orbiculare* actin gene (GenBank accession number, AB778553.1).

### Transformation of BY-2 cells

Tobacco BY-2 cells were grown in Linsmaier and Skoog medium supplemented with 3% sucrose and 0.2 mg/L 2,4-dichlorophenoxyacetic acid at 26°C (48). To generate the *SP*-*SIB1*-expressing transgenic line, pBICHgLBSXVE expressing the artificial transcription factor XVE, which activates transcription by binding with 17β-estradiol (28), and pBICLBSER-ToMV-SP-SIB1 were introduced into tobacco BY-2 cells using the *Agrobacterium* method (49). Transgenic lines were selected on agar medium containing the appropriate selective agents, 50 mg/L hygromycin, 100 mg/L kanamycin, and 500 mg/L carbenicillin. Suspended cells developed from calli were grown in 3 mL of liquid medium in six-well culture plates during the primary screening, after which they were transferred to 150 mL of liquid medium in 500-mL flasks with constant shaking at 135 rpm. After the initial culture for 2–3 weeks, the suspension cells were maintained without selective agents. These cell lines were suspension-cultured in normal MS medium and MS medium labeled with an ^15^N nitrogen source for NMR analysis.

### Protein production and purification

Protein production was induced by adding 10 μM 17β-estradiol (28). After 4 days, SIB1 protein had accumulated in the culture medium, and the culture medium was collected by centrifugation. For the first purification, the ammonium sulfate precipitation method was performed, and the protein in the 60% ammonium sulfate supernatant was mostly SIB1 protein. The solvent of the supernatant was replaced with phosphate buffer (pH 6.8) by dialysis. Next, the supernatant was purified by gel filtration chromatography using AKTA prime plus (GE Healthcare, Buckinghamshire, UK) to obtain a single protein. For the gel filtration chromatography purification, a Superdex 75 10/300 GL column (Amersham Biosciences, Uppsala, Sweden) was used, and the buffer was phosphate buffer (pH 6.8) at a flow rate of 0.1 mL min^−1^ at room temperature. Elution was monitored by absorbance at 280 nm. The collected fraction was concentrated using a centrifugal concentrator (CC-105, Tomy Seiko Inc., Tokyo, Japan) and then used for NMR analysis.

### Gene disruption in *C. orbiculare*

To delete *SIB1*, we first generated the *lig4*Δ strain from 104-T, in which the homologous recombination ratio is expected to be increased, because DNA ligase 4 (Lig4) is reported to be a key molecule in the nonhomologous end-joining pathway (24). To generate the *lig4-*knockout strain, we introduced pBATTEFPGENLIG4KO into protoplasts of *C. orbiculare* 104-T. Preparation of protoplasts and transformation of *C. orbiculare* were performed according to a method described previously (50). We first selected geneticin-resistant transformants, and the bialaphos-sensitive transformants were selected from the geneticin-resistant transformants. The selected bialaphos-sensitive transformants were subjected to genomic PCR analysis using the primers Co5-Jcheck3 and J-check-CoLIG3AS to check the disruption of *LIG4*. The *lig4*Δ strains obtained exhibited colony growth, conidiation, and virulence on cucurbits to the same extent as the parental wild-type strain 104-T. To generate *SIB1-*knockout mutants, we introduced the gene-disruption vector pCB1636SIB1 into protoplasts of the *C. orbiculare lig4*Δ strain (generated in the 104-T background as described above). We selected hygromycin-resistant transformants. Transformants were then analyzed by genomic PCR with the primers SIB1_col_F and SIB1_col_R. Hygromycin-resistant, geneticin-resistant, and bialaphos-sensitive transformants were selected in regeneration medium containing hygromycin B (100 μg/mL), geneticin (200 μg/mL), and bialaphos (25 μg/mL), respectively.

### Inoculation of *N. benthamiana*, cucumber, and melon

Conidial suspensions collected from the 7-day-old colony of each strain formed on PDA were drop-inoculated onto detached *N. benthamiana* leaves, and cotyledons of cucumber and melon; the volume was 10 μL for each drop. All conidial suspensions were used at a concentration of 5 × 10^5^ conidia/mL. In *N. benthamiana*, the leaves were collected from 5–6-week-old plants. The cotyledons of cucumber and melon were derived from 10-day-old plants. The phenotype of lesions developed was observed after incubation for 7 days at 24°C.

### MS analyses

Samples with 0.1% trifluoroacetic acid (TFA) were filtered through a 0.45-μm filter, and the filtrates were injected directly into a C18 column (4.6 mm inner diameter × 250 mm, Protein-R; Nacalai Tesque, Kyoto, Japan) equilibrated with 100% mobile phase A (0.1% TFA in water). Samples were separated with a linear gradient from 0% to 50% mobile phase B (0.1% TFA in acetonitrile) in 40 min at a 0.5-mL/min flow rate. The eluate was monitored at 220 nm, and the fraction including SIB1 was verified by matrix-assisted laser desorption/ionization time-of-flight mass spectrometry (MALDI-TOF-MS; ultrafleXtreme, Bruker, Billerica, MA, USA). The peptide concentration was estimated using a BCA protein assay reagent kit (Thermo Fisher Scientific, Waltham, MA, USA).

To confirm the pyroglutamylation of SIB1, 5 μg (1 μg/μL) of SIB1 dissolved in buffer (6 M urea and 0.1 M triethylammonium bicarbonate (TEAB)) was incubated for reduction with 2 mM Tris(2-carboxyethyl)phosphine (TCEP) for 30 min at 37°C followed by alkylation with 55 mM iodoacetamide (IAA) in the dark for 30 min at room temperature. The sample solution was acidified with 10% TFA and desalted using SDB-Stage Tip (51). The desalted sample was dried under vacuum and dissolved in buffer (50 mM Na_2_PO_4_, pH 7.0, 10 mM dithiothreitol, and 1 mM ethylenediaminetetraacetic acid), and 10 μL (1 mU) of *Pfu* PGAP (TaKaRa) was added. After incubation for 5 h at 50°C, the sample was acidified with 10% TFA, desalted using SDB-Stage Tip, and dried under vacuum. The PGAP-treated sample was analyzed using MALDI-TOF-MS. The Unimod database (http://www.unimod.org) was used to analyze the posttranscriptional modification.

To examine whether Cys residues of SIB1 were in the oxidized form, 15 μg (1 μg/μL) of SIB1 dissolved in buffer (6 M urea and 0.1 M TEAB) was used as the stock solution. Using this stock, three samples at different conditions were prepared: (i) untreated, (ii) alkylated with 55 mM IAA under nonreducing conditions, and (iii) reduced with 2 mM TCEP and alkylated with 55 mM IAA. All three samples were acidified with 10% TFA and desalted using SDB-Stage Tip. The desalted sample was dried under vacuum and analyzed using MALDI-TOF-MS. The disulfide linkages were determined using the method reported previously (52). SIB1 (5 μg, 1 μg/μL) was dissolved in buffer (6 M urea and 0.1 M TEAB) and Lys-C was added to the sample at a 1:100 ratio of Lys-C. After overnight digestion at 37°C, the sample solution was acidified with 10% TFA and desalted using SDB-Stage Tip. The desalted solution was dried under vacuum and analyzed using MALDI-TOF-MS. The assignment of peaks derived from peptides with disulfide linkages was performed using BioTools (Bruker Daltonics).

### NMR study of SIB1

_15_N-labeling of SIB1 using the BY-2 system was performed using the method reported previously (27,29,30), and the sample was purified as described above. The ^15^N-labeled NMR sample was prepared at a concentration of 0.8 mM dissolved in H_2_O containing 10% D_2_O and 100 mM KCl. The sample pH was adjusted to 6.3 by direct reading with a pH meter. All NMR data were recorded on a Bruker AVANCE III 800 equipped with a TCI cryogenic probe. The sample temperature during the NMR experiments was kept at 25.0°C.

To determine the structure, ^1^H-^15^N HSQC (53,54), ^15^N-edited NOESY (55), ^15^N-edited TOCSY (56), NOESY (57) and TOCSY (58) were observed. The NOE mixing time and TOCSY spin-lock time were set as 100 and 70 ms, respectively. In addition, heteronuclear {^1^H}-^15^N NOE experiments (59) were performed to provide information about the internal protein dynamics. Water suppression in the NMR experiments was achieved using WATERGATE (60) or a water flip-back pulse (61). All FID data were processed using NMRPipe (62) and analyzed on Sparky (63). The distance constraints were obtained from NOE peaks. The angle constraints were obtained using TALOS+ (64) analysis using ^1^HN, ^15^N, and α_1_H chemical shifts. The three-dimensional structure of SIB1 was calculated using CYANA (65) and NMR-based constraints. The structural figures were generated using MOLMOL (66) or PyMOL (67).

The pulse sequences used to study protein dynamics have been published (68). In our study, the *T*_1_ relaxation analysis used a series of 10 experiments with relaxation delays set at 75, 100, 150, 200, 250, 300, 400, 500, 700, and 950 ms. Similar to the *T*_1_ experiments, *T*_2_ measurements were also performed as a series of 10 experiments with different relaxation delays of 25, 60, 80, 120, 150, 200, 300, 400, 500, and 750 ms. The *T*_1_ and *T*_2_ values were estimated by fitting the peak volume, *I*, using the equation, *I* = *I*_o_ exp(–*t*/*T*_1,2_). As for the heteronuclear NOE experiments, a 5-s recycle delay was used after each scan. The NOE values were obtained by calculating the ratio of the peak intensity recorded with the saturation of protons divided by the peak intensity recorded without saturation.

## Data deposition

Protein structure coordinate data are available at Protein Data Bank (PDB) (https://www.rcsb.org/). Accession codes for the structural coordinates and chemical shifts deposited in the PDB and Biological Magnetic Resonance Data Bank (BMRB) are 7EAU and 36412, respectively. RNAseq data are accessible in NCBI’s Gene Expression Omnibus (GEO) database under GEO Series accession number GSE178879.

## Acknowledgements

The authors thank all technicians working at Center for Nano Materials and Technology (CNMT), Japan Advanced Institute of Science and Technology (JAIST) for maintenance of instruments used in this work. The authors also thank Akiko Mizuno and Hiroko Hayashi in Ishikawa Prefectural University and Fumika Takahashi in Kyoto University for excellent technical assistance. The authors also thank Manaka Iino for his contribution to a part of the NMR data analyses. This work was supported by Grants-in-Aid for Scientific Research (18H02204 and 21H04725 to Y.T., 17K08194 to S.O., 17H06172 to K.S., 19K15846 to P.G) (KAKENHI).

## Abbreviations

BY-2: *Nicotiana tabacum* cv. Bright Yellow 2
EK: enterokinase
ETI: effector-triggered immunity
HR: hypersensitive response
HSQC: heteronuclear single quantum coherence spectroscopy
IAA: iodoacetamide
MALDI-TOF-MS: matrix-assisted laser desorption/ionization time-of-flight mass spectrometry
NMR: nuclear magnetic resonance
NOESY: nuclear Overhauser effect spectroscopy
PAMP: pathogen-associated molecular patterns
PCD: programmed cell death
PTI: PAMP-triggered immunity
RBOHs: respiratory burst oxidase homologs
ROS: reactive oxygen species
SDS-PAGE: sodium dodecyl sulfate–polyacrylamide gel electrophoresis
TCEP: Tris(2-carboxyethyl)phosphine
TEAB: triethylammonium bicarbonate
TFA: trifluoroacetic acid
TOCSY: total correlation spectroscopy

